# Extrachromosomal DNA in the cancerous transformation of Barrett’s esophagus

**DOI:** 10.1101/2022.07.25.501144

**Authors:** Jens Luebeck, Alvin Wei Tian Ng, Patricia C. Galipeau, Xiaohong Li, Carissa A. Sanchez, Annalise Katz-Summercorn, Hoon Kim, Sriganesh Jammula, Yudou He, Scott M. Lippman, Roel Verhaak, Carlo C. Maley, Ludmil B. Alexandrov, Brian J. Reid, Rebecca C. Fitzgerald, Thomas G. Paulson, Howard Y. Chang, Sihan Wu, Vineet Bafna, Paul S. Mischel

## Abstract

**BACKGROUND:** Oncogenes are commonly amplified on extrachromosomal DNA (ecDNA) contributing to poor outcomes for patients. Currently, the chronology of ecDNA development is not known. We studied the origination and evolution of ecDNA in patients with Barrett’s esophagus (BE) who progressed to esophageal adenocarcinoma (EAC).

**METHODS:** We analyzed whole-genome sequencing (WGS) data from a BE surveillance cohort and EAC patients at Cambridge University UK (n=206 patients). We also analyzed WGS data from biopsies taken at two time points from multiple sites in the esophagus from 80 patients enrolled in a case-control study at the Fred Hutchinson Cancer Center (FHCC) - 40 BE patients who progressed to EAC and 40 who did not.

**RESULTS:** ecDNA was detected in 24% and 43% of BE patients with BE-associated early and late-stage EAC, respectively, in the Cambridge cross-sectional cohort. ecDNA was found in 33% of all FHCC BE patients who developed cancer, either prior to, or at EAC diagnosis. ecDNA was strongly associated with patients who developed cancer, in contrast with FHCC BE patients who did not progress (odds ratio, 18.8, CI – 2.3-152, p=3.3×10-4). ecDNAs were enriched for oncogenes and immunomodulatory genes and could be detected early in the transition from high-grade dysplasia to cancer and increased in copy number and complexity over time.

**CONCLUSIONS:** ecDNAs can develop before a diagnosis of cancer in BE patients and are strongly selected for during the evolution to EAC. ecDNAs promote diverse oncogene and immunomodulatory gene amplification during EAC development and progression.

## INTRODUCTION

EAC is a highly lethal cancer that can arise from BE, a relatively common, pre-cancerous metaplastic condition (1.6% of the U.S. population)^1^. In addition to epidemiological and clinical features such as chronic gastroesophageal reflux disease, patient age or BE lesion size^2,3^, genomic copy number changes have also been implicated in the transformation to EAC^1,4–9^. Such changes include oncogene amplifications, which frequently occur on circular extrachromosomal DNA (ecDNA) particles^10^. The non-chromosomal inheritance and highly accessible chromatin architecture of ecDNA contributes to aggressive tumor growth, accelerated evolution, and drug-resistance^11–14^. Computational tools can detect ecDNA in sequencing (WGS) data from biopsies^15–17^. However, the relative lack of pre-cancer to cancer longitudinal studies, plus the challenges of interpreting clonality in the face of non-Mendelian genetics, have made it difficult to determine if ecDNAs arise early in tumorigenesis and contribute to the transformation of dysplasia into cancer. Two carefully curated surveillance studies of BE patients, including a longitudinal case-control study with multi-regional WGS sampling, and a completely independent cross-sectional surveillance cohort, with full histological correlatives, provided an unprecedented opportunity to study the role of ecDNA in the transition of BE to EAC.

## METHODS

### Study samples

We analyzed WGS data from 206 patients in a cross-sectional BE surveillance Cambridge cohort with biopsy-validated BE, including 42 patients with metaplasia or low-grade dysplasia (LGD) who never developed HGD or EAC during follow-up, 25 patients with high-grade dysplasia (HGD), 51 patients with early-stage EAC (AJCC stage I), and 88 patients with late-stage EAC (AJCC stage II-IV) (Figure 1a). Histology and WGS sequencing were performed on the same biopsies (Supplementary Appendix – “Cambridge sample selection”). We also analyzed 20 EAC tumors from The Cancer Genome Atlas (TCGA) esophageal carcinoma study^18^ (ESCA), composed of 6 early-stage and 14 late-stage tumors.

**Figure 1.**
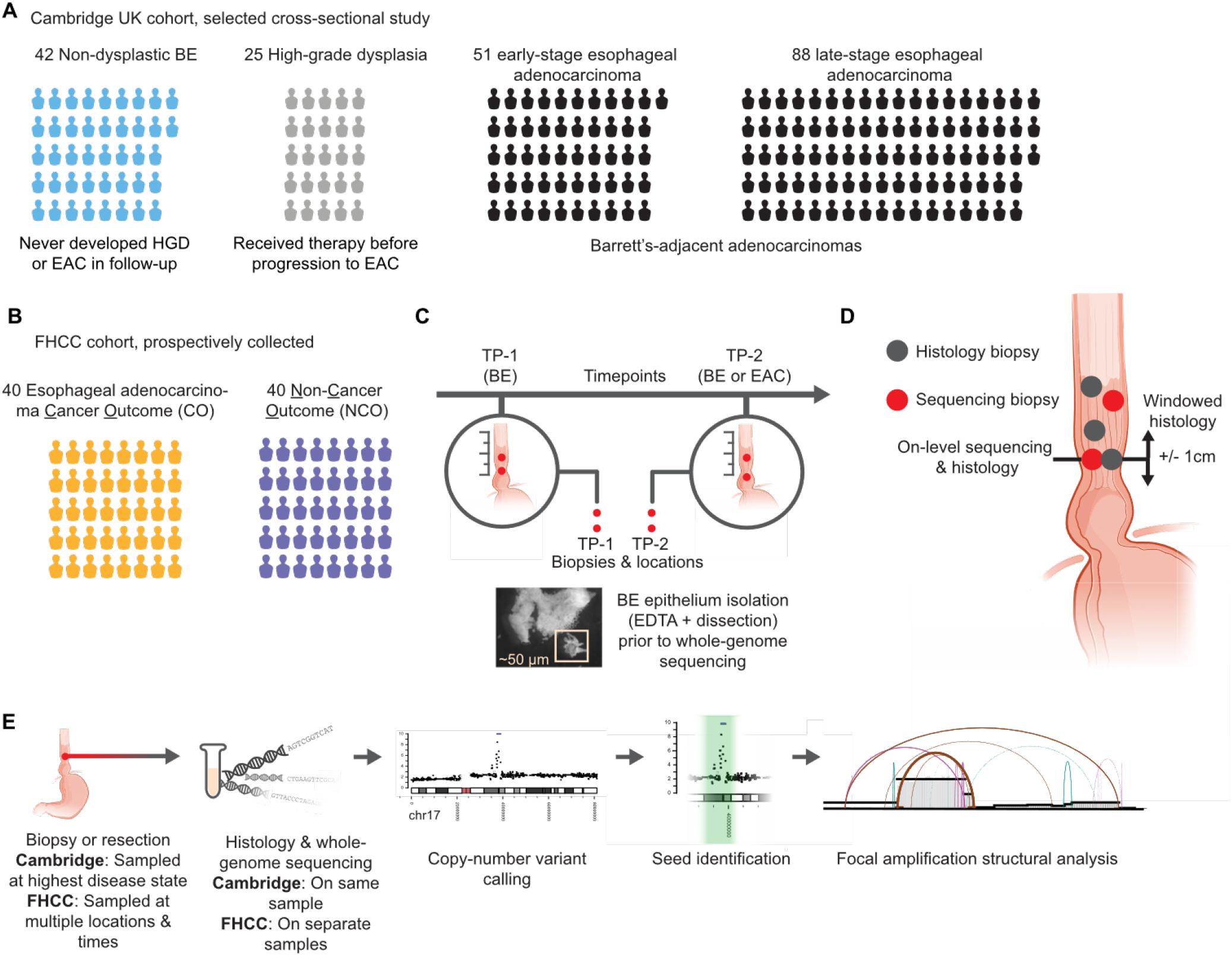
Study designs. **a)** Breakdown of the types of BE patients in the Cambridge cross-sectional study, segregated into cohorts by histology of the sample sequenced, which was also the highest-disease state for that patient. **b)** The FHCC cohort consists of 80 patients for whom biopsies were collected prospectively and later separated in two groups of 40 patients who had cancer outcomes (CO) and non-cancer outcomes (NCO) **c)** Sample collection at timepoints TP-1 and TP-2 to collect sequencing biopsies and histology biopsies. Two sequencing biopsies were collected from each timepoint. **d)** WGS biopsies and histology biopsies were taken independently. Some histology and sequencing biopsies were taken at the same level of the esophagus (on-level), and some histology biopsies fell within a +/- 1cm window of the measured height of the sequencing biopsy (windowed histology). **e)** Workflow for analyzing the WGS samples. A brief overview of the process by which biopsies were selected, sequenced, and characterized by AmpliconArchitect is shown.

We analyzed WGS data from esophageal biopsies in an independent, prospectively collected case-control study conducted at the Fred Hutchinson Cancer Center (FHCC) of patients with BE (Figure 1b.)^8^, including 40 patients with a cancer outcome (CO) and 40 patients who did not develop cancer (NCO) during the study period. At least two biopsies for WGS were obtained by isolating epithelial tissue (Figure 1c) of the BE at each of the two primary study time points - timepoint 1 (TP-1) and timepoint 2 (TP-2). Histology samples were also collected independently of the sequencing biopsies, including from the same, or close to the same, level in the esophagus (Figure 1d). At TP-2, biopsies from the same level of the esophagus (or as close as possible) as the EAC were used for sequencing (Supplementary Appendix), except for patient 391 for whom resected tumor sequencing was available.

### ecDNA detection and characterization

DNA copy number alterations were detected using CNVKit^19^ (FHCC, TCGA cohorts) and ASCAT^20^ (Cambridge cohort) with PrepareAA (https://github.com/jluebeck/PrepareAA) to identify candidate seed regions for ecDNA detection and characterization using AmpliconArchitect^15^ (AA) and AmpliconClassifier (Supplementary Appendix – “Amplicon classification and ecDNA detection”) (Figure 1e). An amplicon complexity score was computed based on the diversity of amplicon structure decompositions output by AmpliconArchitect (Supplementary Appendix - “Amplicon complexity score”).

### Statistical analysis

We used SciPy^21^ (version 0.19.1) to conduct all statistical tests in the study, with the exception of the ecDNA region/oncogene overlap significance test which utilized ISTAT^22^ (version 1.0.0). When computing odds ratios, if any cell in the two-by-two table was zero, the Haldane correction^23^ was applied to every cell in the table. Significance of odds ratios and differences in event frequencies between groups were assessed by Fisher’s exact test. The default alternate hypothesis used was “two-sided” in relevant statistical tests, unless otherwise specified as H_a_=greater or less.

## RESULTS

### ecDNA is significantly associated with EAC in patients with BE

ecDNA was not detected in any of the non-dysplastic Barrett’s esophagus (NDBE) samples in the cross-sectional BE surveillance Cambridge cohort (Supplemental Figure 1). By contrast, ecDNA was found in tumors from 13/51 patients (25%) with early-stage (AJCC stage I) EAC and in tumors from 38/88 patients (43%) with late-stage tumors (AJCC stage II-IV) (Figure 2a). ecDNA occurrence was significantly enriched in early-stage EAC versus NDBE (Figure 2b, p=1.8×10^−4^) with significantly increased ecDNA frequency in late-stage tumors compared to early-stage (odds ratio (OR)=2.2, C.I.=1.0-4.7, Fisher’s exact test, p=0.027, Ha=greater). ecDNA was detected in a nearly identical fraction of an independent cohort of late-stage EAC tumors from TCGA (6/14 tumors – 43%).

**Figure 2.**
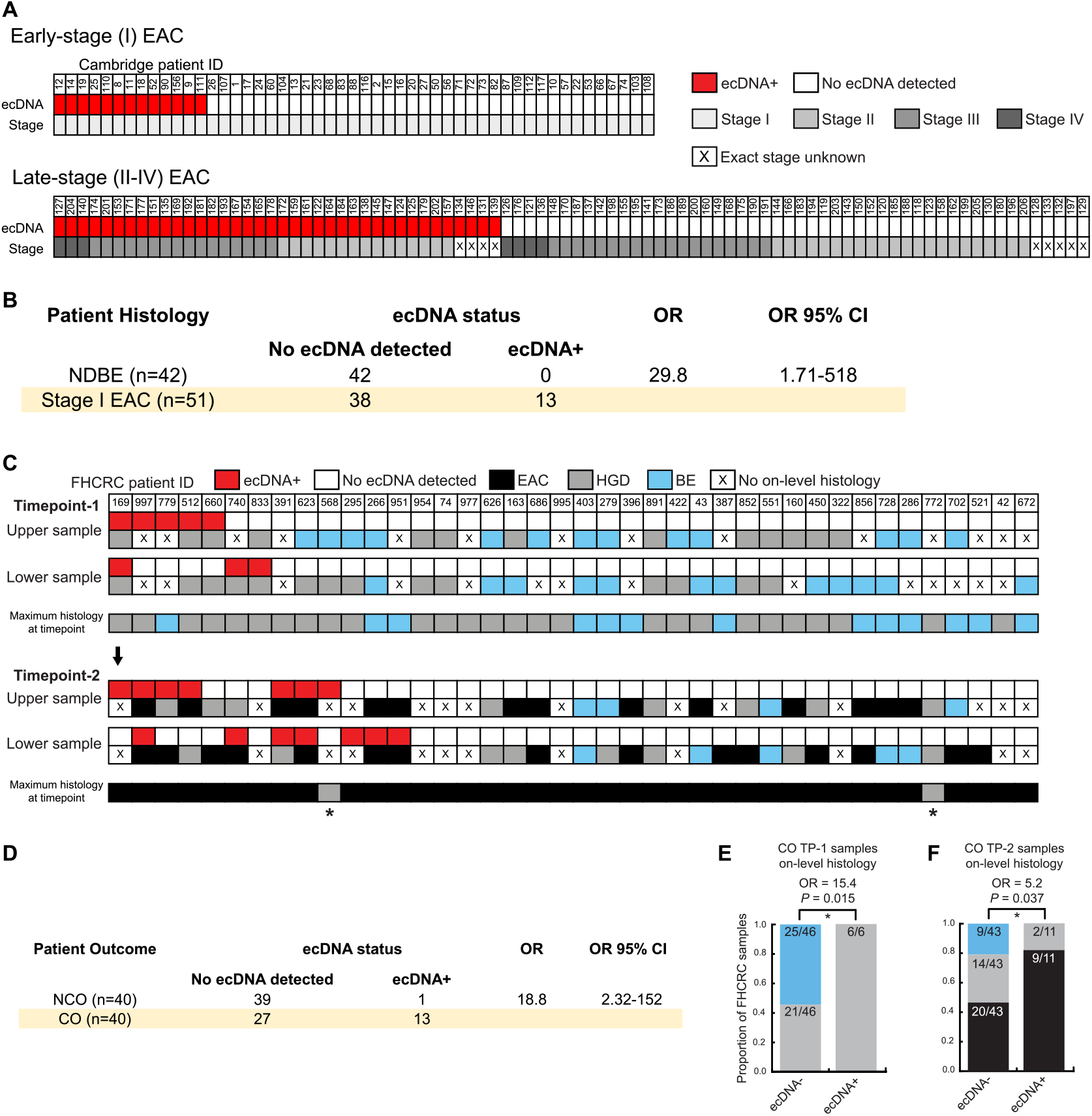
**a)** Characterization of the ecDNA status and cancer stage of patient samples from the Cambridge early- and late-stage EAC cohorts. **b)** Comparison of patient ecDNA status and cancer outcome status reveals association of ecDNA with EAC (Fisher’s exact test, p=1.8×10^−4^, Ha=greater). **c)** Characterization of the ecDNA status and on-level histology of samples collected for FHCC cancer outcome (CO) patients across timepoints TP-1 and TP-2 for the two esophageal sequencing samples (“upper” and “lower”). Maximum histology of any biopsy from that timepoint is also shown. Asterisk indicates cancer diagnosis made at next endoscopy (1.44 and 8.16 months after TP-2 for patients 568 and 772, respectively). **d)** Comparison of patient ecDNA status and cancer outcome status reveals association of ecDNA with cancer outcome (odds ratio = 18.8, C.I. = 2.3-152, Fisher’s exact test, p=3.3×10^− 4^, Ha=greater). **e)** The proportion of FHCC TP-1 samples without EAC or HGD in on-level histology, versus with HGD in on-level histology for CO patients (before developing cancer), segregated by ecDNA status, shows enrichment for ecDNA with advanced disease status (OR=15.4, Fisher’s exact test, p-value=0.015, H_a_=greater). **f)** The proportion of FHCC TP-2 samples without HGD or EAC in on-level histology (BE only) versus with HGD or EAC in on-level histology in CO patients (cancer first detected), segregated by ecDNA status shows association with HGD/EAC status OR=5.2, Fisher’s exact test, p-value=0.037, Ha=greater.

The FHCC study featured multi-regional, longitudinal sampling from before and at cancer diagnosis. We examined the development of ecDNA over time in patients who progressed to EAC endpoint versus those who maintained benign BE. At cancer diagnosis (TP-2), ecDNA was detected in samples from 11/40 CO patients with EAC (28%) (Figure 2c, Supplemental Figure 2), consistent with the 25% ecDNA frequency found in the Cambridge cohort of patients with early-stage cancer (Fisher’s exact test, p=1.0). ecDNA was detected in only 1/40 non-cancer outcome (NCO) patients (Supplemental Figure 3). Notably for this sample, KRAS was amplified (Supplementary Figure 4), and the patient died of causes unrelated to BE only 2.84 years after TP-2. We additionally analyzed 20 long-term follow-up samples collected from 10 NCO patients (median 9.6 years after TP-2) who maintained NDBE status and remained ecDNA-negative (Supplemental Figure 5). These data demonstrate a highly significant association of ecDNA with the development of EAC (Figure 2d).

**Figure 3.**
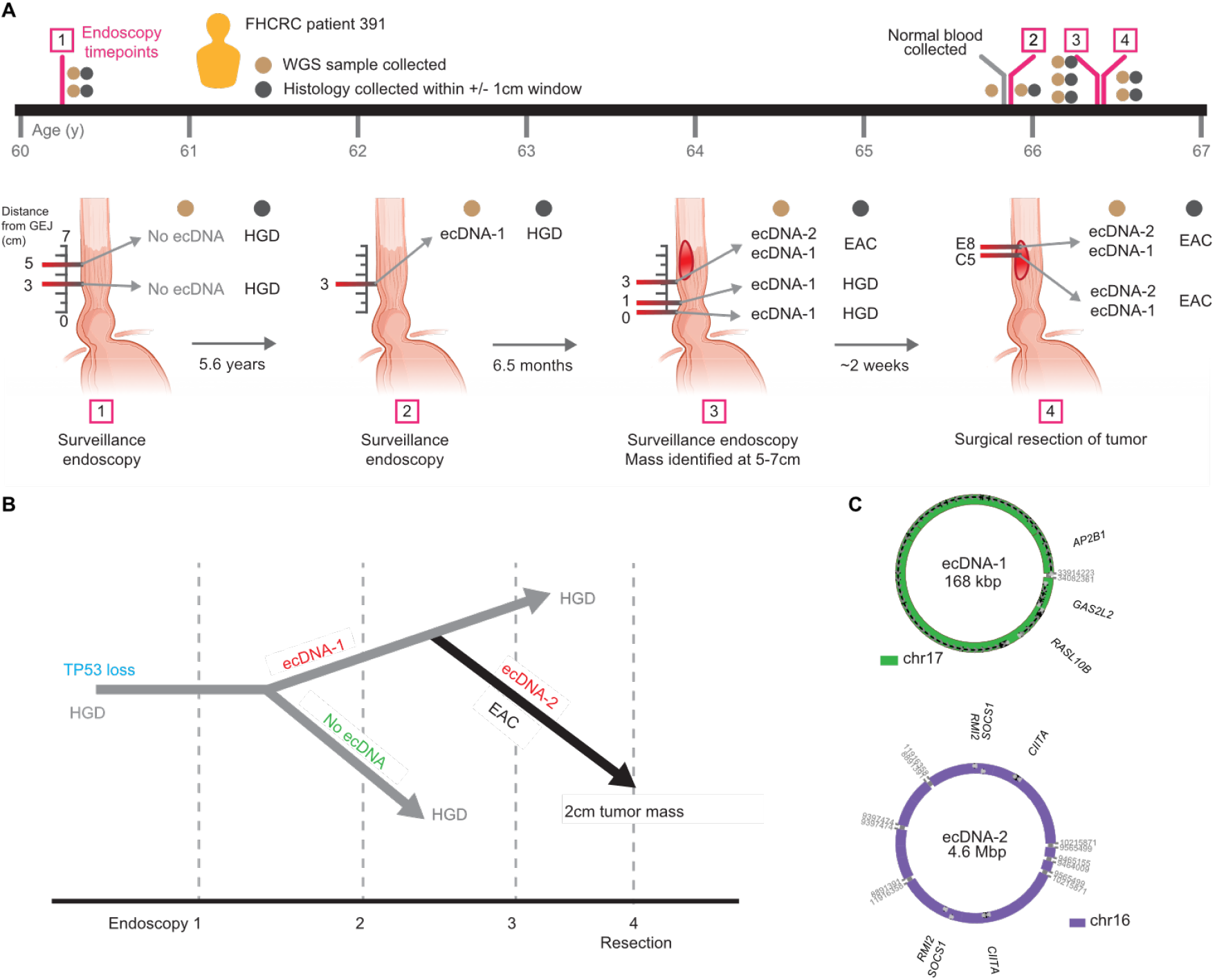
**a)** Timeline of sample collection in FHCC CO patient 391 relative to patient age. Summary of the ecDNA status and windowed histology status for four endoscopies with time interval between each also indicated. Biopsy distances from the gastroesophageal junctions (GEJ) are indicated. Two distinct species of ecDNA are labeled as ecDNA-1 and ecDNA-2. **b)** Inferred phylogeny of patient 391 WGS samples across the four endoscopies, starting from *TP53* loss, with branching reporting the ecDNA formation events, annotated by histological status of sample (windowed). **c)** The structure of ecDNA-1, first detected in endoscopy-2 where HGD was detected within +/- 1cm, and the structure of ecDNA-2, first detected in endoscopy-3 where EAC was diagnosed and present within +/- 1cm.

### ecDNA can be detected in esophageal biopsies associated with high-grade dysplasia

The longitudinal case-control FHCC study enabled determination of the timing of ecDNA development in BE patients with a cancer outcome. Remarkably, ecDNA was found at TP-1, prior to development of cancer, in biopsy tissues from 7/40 CO patients (18%) who subsequently developed EAC.. Also, at TP-1, HGD was detected in at least one histology biopsy for 27/40 patients (67.5%). Six TP-1 samples having ecDNA could be matched to an on-level histology biopsy, all showing HGD (Figure 2e). By contrast, 46% (21/46) of the ecDNA-negative TP-1 sequencing biopsies could be matched to on-level HGD, indicating a significant association of ecDNA and HGD in the pre-cancer samples (p-value=0.015, Figure 2e).

In CO patient samples collected at TP-2, where cancer was first diagnosed, we associated 54 sequencing biopsies to on-level histology. ecDNAs were identified in 11 of these sequencing biopsies, nine of which (82%) associated with on-level EAC, with the remaining 2/11 associated with on-level HGD (Figure 2f). In contrast, among the remaining 43 ecDNA-negative biopsies, only 20/43 (47%) were associated with on-level EAC, with the remaining 23/43 (53%) associated with on-level BE or HGD (Figure 2f). The specificity of ecDNA association with worsened pathological status at both timepoints suggests that ecDNA are enriched in BE clones that become cancer. In the Cambridge cohort, ecDNA was detected in only one of the 25 patients with HGD (Supplemental Figure 1). However, in that cohort, HGD was treated immediately upon detection, so it was not possible to determine if the HGD samples would have subsequently progressed to cancer.

### *TP53* loss and ecDNA formation

Prior loss of *TP53* enables genomic instability^7,8,24,25^, and we found a strong association in both FHCC and Cambridge cohorts between *TP53* disruption (Supplementary Appendix) and ecDNA-positive status (Supplemental Figure 6a-b). In the FHCC cohort, all eight samples in which ecDNA was found prior to cancer diagnosis (TP-1) showed biallelic disruption of *TP53*. The appearance of ecDNA as a subset of *TP53* disrupted cases points to the prior loss of *TP53* enabling ecDNA formation.

### Ongoing ecDNA formation associates with the malignant transformation

To better understand the potential relationship between ecDNA and the transition from HGD to EAC, we studied an individual patient in the FHCC CO cohort (patient 391) for whom WGS data were collected at four endoscopies over a seven-year period (Figure 3a). Initially, HGD was detected at two different locations within the BE segment. Chromosomal *ERBB2* amplification via breakage fusion bridge cycles (Supplemental Figure 7a) and *TP53* loss were present in these biopsies (Figure 3b). An ecDNA (ecDNA-1), bearing *AP2B1, GAS2L2* and *RASL10B*, was only detected (Figure 3b-c, Supplemental Figure 7b) after 5.6 years. The lesions did not progress to EAC for another 6.5 months, at which point a second ecDNA (ecDNA-2) containing *SOCS1, CIITA*, and *RMI2* was detected (Figure 3b-c, Supplemental Figure 7c). *SOCS1* is a suppressor of cytokine signaling, including interferon gamma^26^ that may foster escape from cytotoxic T-cells^27^. *CIITA* is an immunomodulatory master transcription factor for antigen presentation^28^, whose translocation is immunosuppressive^29^. *RMI2* is a component of the Bloom Helicase complex involved in homologous recombination that has been suggested to play a role in lung cancer metastases^30^. A subsequent surgical resection of the tumor confirmed both ecDNA-1 and ecDNA-2, whereas the tissue containing only ecDNA-1, *TP53* loss and chromosomal *ERBB2* amplification, remained HGD. These results suggest that multiple and ongoing focal amplification events occur in dysplastic tissues^31,32^, enhancing a clone’s fitness during malignant transformation.

### Genomically overlapping ecDNAs reidentified at multiple timepoints share a common origin

To compare ecDNA fine structure across multiple time points in the same individual, we developed an amplicon similarity score ranging from 0 to 1 (Supplemental Figure 8a-e, Supplementary Appendix – “Amplicon similarity score”). Genomically overlapping ecDNAs from multiple samples from the same patient shared high similarity, consistent with a common origin (Supplemental Figure 8e). All ten genomically overlapping ecDNA pairs from within the same FHCC patients reidentified between TP-1 and TP-2 showed significant similarity (p<0.05) (Supplemental Table 2). Thus, ecDNAs detected in pre-cancer are frequently maintained through the transition to cancer, and genomically overlapping ecDNA identified from multi-region sampling likely have a common origin. Taken together, these data suggest that ecDNA can be a truncal event in the formation and evolution of EAC.

### ecDNAs are selected for and evolve during the transformation of BE to EAC

We detected a marked increase in ecDNA frequency in cancer-outcome patients prior to clinical detection of cancer, and even more elevated levels in later-stage cancers (Figure 4a). To better understand these observations, we characterized 137 ecDNA across all samples from 75 ecDNA-positive BE, BE-derived HGD, and BE-adjacent EAC patients. ecDNA copy number was significantly higher in EAC samples than in pre-cancer samples (Figure 4b). Moreover, the complexity of structural rearrangements in ecDNA-derived regions increased between pre-cancer and EAC (Figure 4c), suggesting a significant increase in the heterogeneity of ecDNA structures with the evolution of tumors. We next investigated CN changes in 8 pairs of clonal ecDNA where the same ecDNAs (based on amplicon similarity score) reappeared in different sequencing samples from the same patient, and for which the samples had windowed histology data available (Supplementary Table 2). When both ecDNA occurrences were associated with the same histology, the ecDNA CNs were highly similar. However, if one sample associated with a more severe histological status than the other, the ecDNA copy number was significantly higher in that sample (Figure 4d). These data suggest that ecDNA confer a strong selective advantage to the BE clones that eventually progress to EAC, and pre-cancer ecDNA are subject to continued evolution during malignant transformation and progression, leading to increased heterogeneity and copy number.

**Figure 4.**
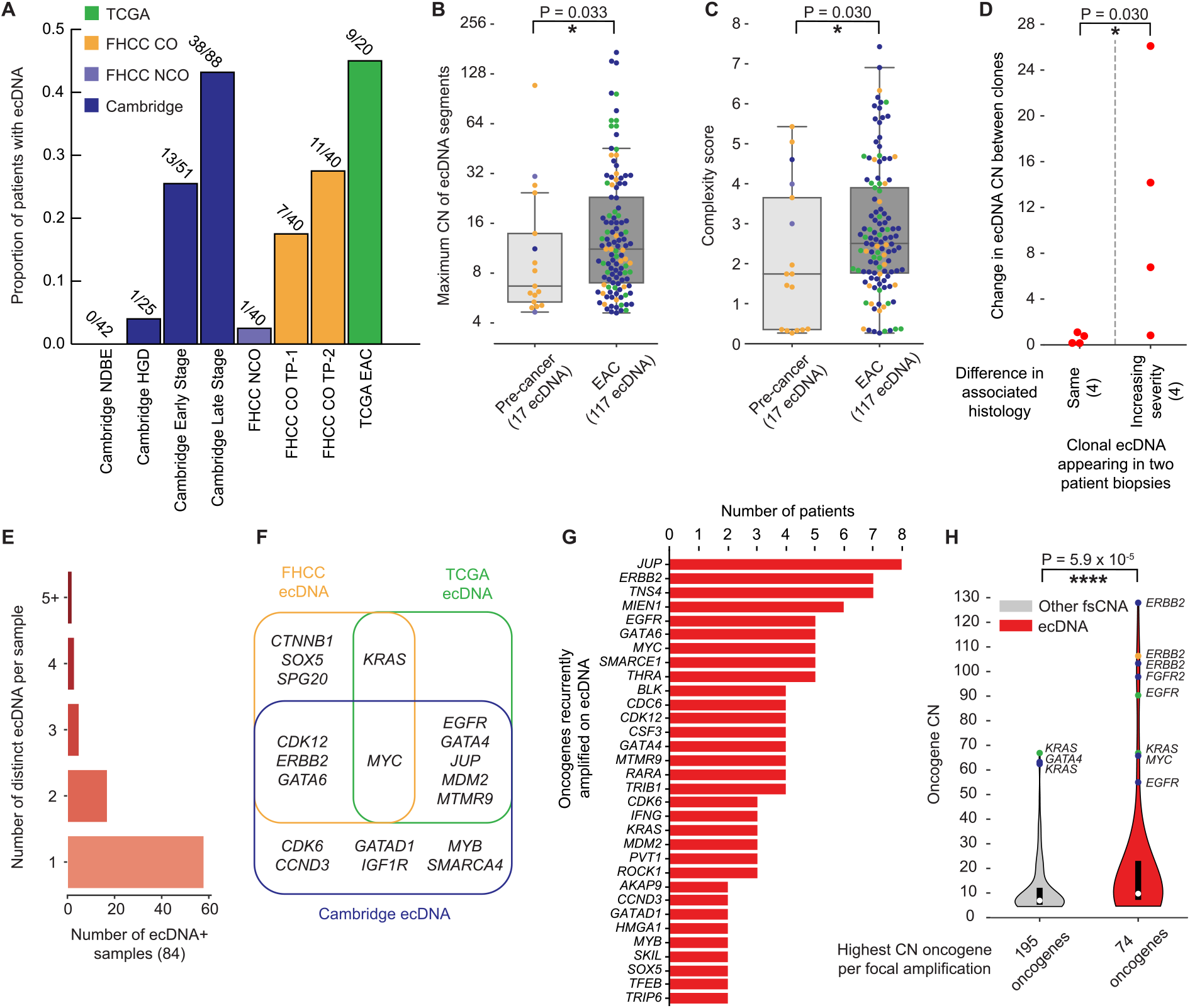
**a)** The proportion of patients with ecDNA in all study cohorts. **b)** The maximum genomic copy number of each ecDNA in pre-cancer samples and EAC (or EAC-linked for FHCC) samples, colored by sample study source. **c)** The complexity score of focally amplified ecDNA-positive genomic regions for pre-cancer and EAC samples. **d)** For clonal ecDNA identified across multiple FHCC samples (determined by amplicon similarity scoring), the increase in ecDNA copy number for each pair of clonal ecDNA, separated by difference in associated histology of the two samples – showing an association of increasing copy number with increasing histological severity (Mann-Whitney U test, p-value=0.030, test statistic=1.0). **e**) The number of distinct ecDNA per sample identified in ecDNA-positive samples from all combined sources of data. **f)** Comparative overlap of BE-associated oncogenes found on ecDNA in the four cohorts. **g)** For oncogenes recurrently detected on ecDNA in different patients, the number of ecDNA-positive patients having the oncogene listed on ecDNA. **h)** Oncogene copy number for the highest copy number focally amplified oncogene on each unique focal amplification (ecDNA or non-ecDNA fsCNA) shows significantly higher oncogene copy number on ecDNA versus non-ecDNA fsCNA (Mann-Whitney U test, p-value=5.9×10^−5^, test statistic=5020.5).

26/83 (31%) of combined ecDNA-positive samples with definite or associated histology contained more than one species of ecDNA (Figure 4e), enabling multiple oncogene amplifications. However, there were no significant differences between pre-cancer (3/14, 21% having multiple ecDNA species) and EAC samples (23/69, 33% having multiple ecDNA species) (p-value=0.53), suggesting that tumors may achieve subclonal ecDNA heterogeneity early on, and that competition between multiple distinct ecDNAs may play a role in EAC evolution.

### ecDNAs promote diverse oncogene and immunomodulatory gene amplifications during EAC development and progression

Oncogenes known to drive EAC, including *ERBB2, KRAS*, and *MYC*^33,34^, were recurrently detected on ecDNAs found in BE and EAC across multiple cohorts (Figure 4f-g; Supplemental Table 3), suggesting that distinct ecDNA amplify similar oncogenes. Furthermore, many ecDNA-borne oncogenes were not detected on focal, non-ecDNA amplifications (Supplemental Figure 10). ecDNA carried 0.76 unique oncogenes per amplicon (97 oncogenes in 127 ecDNA), compared to 0.52 (192/373) unique oncogenes per amplicon in non-extrachromosomal focal somatic copy number amplifications (fsCNAs), suggesting ecDNA may permit a wider variety of oncogene amplifications.

ecDNA amplification was associated with a greater maximum oncogene copy number than other fsCNAs (distribution mean CN 11.6 and 21.3 for non-ecDNA and ecDNA respectively) (Figure 4h), with some ecDNA genes surpassing 100 copies. ecDNA also permitted greater diversity in maximum CN than non-ecDNA fsCNA (CN variance = 687.9 versus 122.2 in non-ecDNA fsCNA, p-value=1.5×10^−4^). Notably, many (79 total) ecDNA genes were associated with immunomodulation^35^ (Supplemental Table 4, Supplemental Figure 11a), with only 25 of the 79 already present in the set of canonical oncogenes. The ecDNA-amplified immunomodulatory genes achieved a significantly higher CN compared to those on other fsCNAs (p-value=4.1×10^−3^, Supplemental Figure 11b).

Comparing genomic regions predicted to be on ecDNA to oncogene intervals known to associate specifically with BE and EAC^8,33,34^ (Supplemental Table 5), demonstrated a statistically significant overlap (p=3.1×10^−5^, Supplemental Figure 12), suggesting that, despite the high diversity of ecDNA-borne oncogenes, ecDNAs are positively selected in a manner specific to cancer type.

## DISCUSSION

Oncogene amplification on ecDNA enables tumors to evolve at an accelerated rate, driving rapid therapeutic resistance and contributing to shorter survival for patients^10,13,36^. It has been unclear whether ecDNA can contribute to the transformation of pre-cancer to cancer, or whether it is a later manifestation of tumor genome instability. In multiple cohorts of BE patients, we demonstrate that ecDNA appear in HGD, and their presence is strongly associated with EAC progression.

Typical phylogenetic approaches to track cancer clonality assume chromosomal inheritance. Consequently, it has been challenging to infer the clonality and evolution of ecDNA-driven cancers. Our results demonstrate that in tumor evolution from pre-cancer to cancer, ecDNA confers a strong selective advantage to the BE clones that eventually progress to EAC. The remarkable heterogeneity in ecDNA-containing cancers may promote rapid and frequent branching of the phylogenetic tree, fostered by the non-chromosomal inheritance of ecDNA during cell division. Further, the increased prevalence and complexity of ecDNA structures in esophageal cancer samples suggests ongoing selection and evolution during tumor formation and progression^37^.

Our results strongly suggest that ecDNAs usually arise in regions of HGD in BE patients, and nearly always in the context of *TP53* loss. These results complement the recent finding that *TP53* alteration and altered copy number may drive the transition from metaplasia to dysplasia^4,7,8,38^, showing the cooperative nature of various genetic and epigenetic alterations, and suggesting that ecDNA formation may represent a particularly potent driver of transformation and an opportunity for specific therapeutic intervention.

Freed from Mendelian constraints, ecDNA amplifies a broader range of oncogenes, and their copy numbers increase rapidly and dramatically in EAC, consistent with strong positive selection. Increased ecDNA heterogeneity may also foster adaptation to changing conditions. Importantly, the clonal selection and maintenance of immunomodulatory genes on ecDNA prior to cancer development may aid in immune evasion. Taken together, these results raise the possibility that ecDNA may contribute to the development of cancer through multiple mechanisms.

## ACKNOWLEDGEMENTS

The eDyNAmiC research was facilitated by Cancer Grand Challenges (grant ref CGCSDF-2021\100007) with support from Cancer Research UK and the National Cancer Institute. This study was supported by a grant from The National Brain Tumour Society and NIH R01-CA238349 to P.S.M. Research conducted in the Bafna lab (V.B., J.L.) was supported in part by grants U24CA264379, R01GM114362, OT2CA278635 from the NIH, and CGCATF-2021/100025 from CRUK. S.W. is a scholar of and is supported by the Cancer Prevention and Research Institute of Texas (RR210034). Research conducted at Fred Hutchinson Cancer Center (T.G.P., B.J.R., C.A.S., P.C.G., X.L.) was provided by NIH grants P01 CA91955 and P30 CA015704. A.C.K.-S. was funded by Cancer Research UK via a clinical research fellowship. H.K. is supported by Brain Korea 21 Four Project, the Korean Ministry of Food and Drug Safety grant (21153MFDS607), and the Korean Ministry of Science and ICT grant (NRF-2019R1A5A2027340). R.G.W.V. acknowledges support by NIH/NCI grants R01 CA237208, R21 CA256575, R33 CA236681 and Cancer Center Support Grant P30 CA034196, as well as NIH/NINDS grant R21 NS114873. Work in the Alexandrov lab (L.B.A., Y.H.) was supported by the US National Institute of Health’s grants R01ES030993-01A1 and R01ES032547-01 and a Cancer Grand Challenges Mutographs team award funded by Cancer Research UK [C98/A24032]. LBA is also personally supported by a Packard Fellowship for Science and Engineering. This work was also supported by the Pre-Cancer Genome Atlas (PCGA) project with core funding from UC San Diego NCI P30 (P30 CA023100; SML), and in part by the SU2C-AACR-DT25-17, and US National Institute of Health’s grants R01DE026644, P30 CA023100, HHSN261201200031I, UG1CA242596. Ginny Devonshire ran the Cambridge sequencing data analysis pipeline. The esophagus illustration shown in our study was created by scientific illustrator Tami Tolpa (Tolpa Studios, www.tolpa.com).

---

**Supplemental Figure 1.**
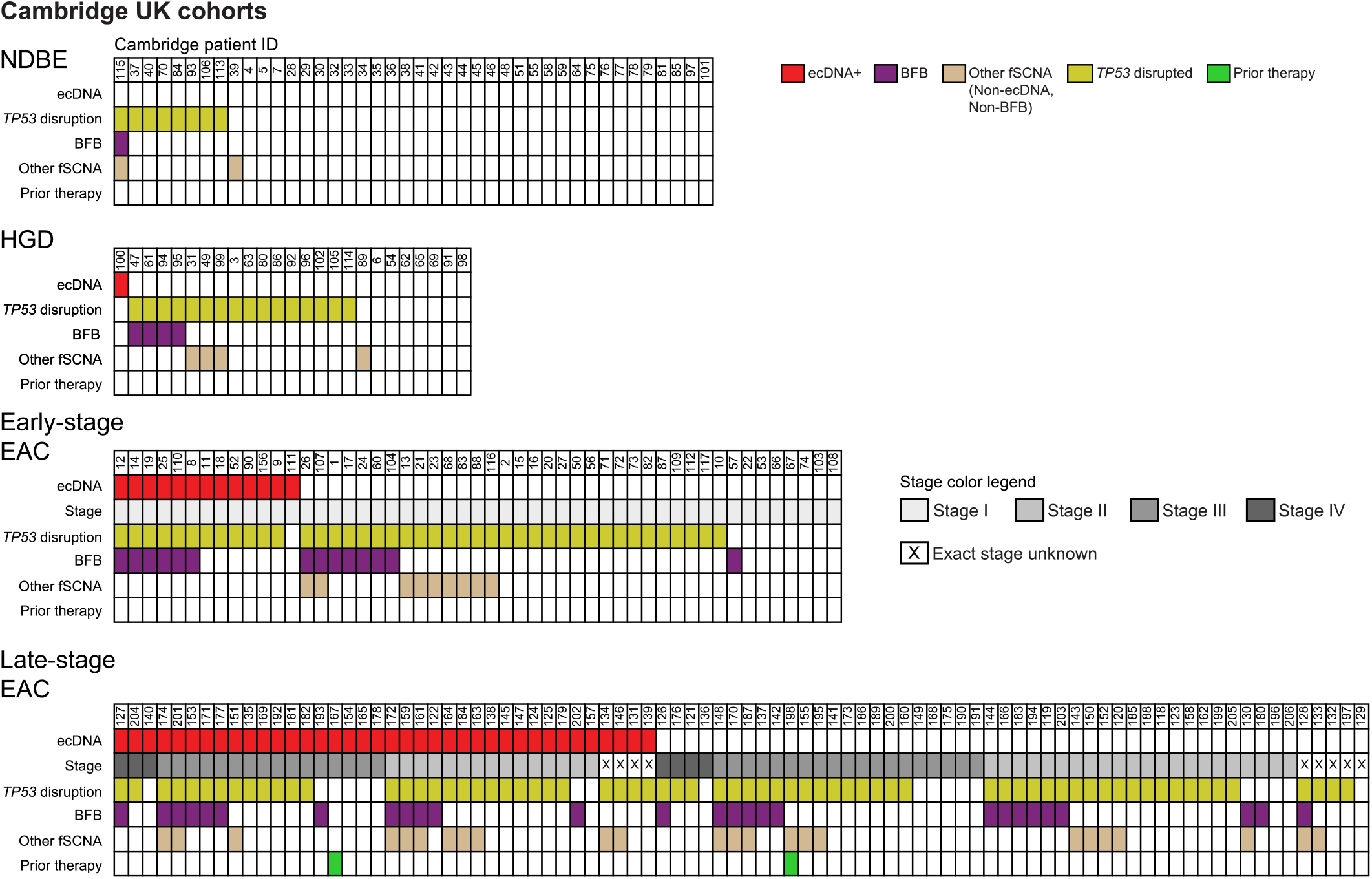
Oncoprint table for Cambridge BE and EAC patients segregated by histology type showing ecDNA status, cancer stage (if applicable) *TP53* disruption (via mutational analysis, involving at least one copy), BFB status, other fsCNA (non-BFB, non-ecDNA) status, and prior therapy (chemotherapy or radiation) on the tumors in cancer patients.

**Supplemental Figure 2.**
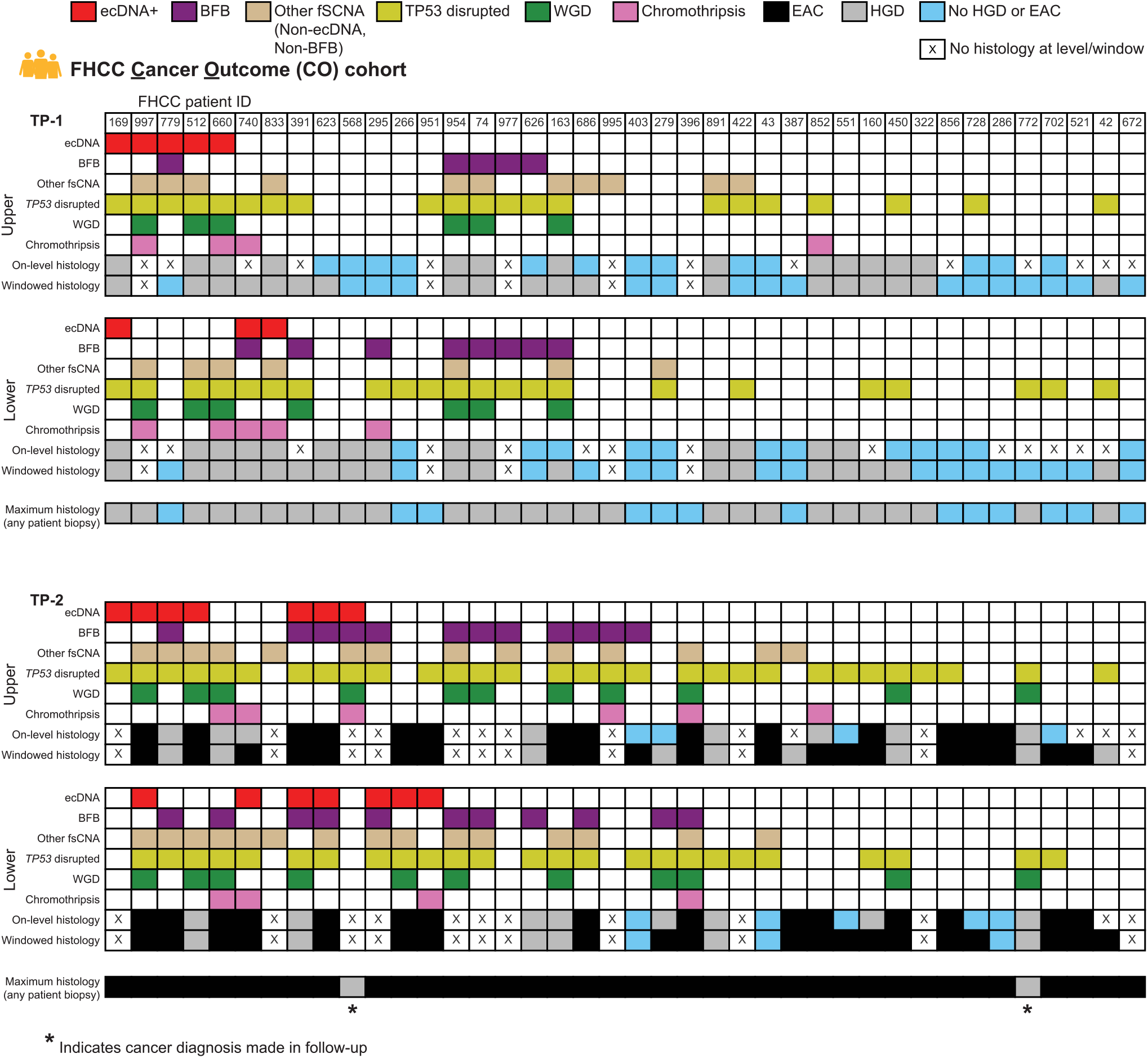
Oncoprint tables of FHCC CO patient WGS samples encoding ecDNA status, BFB status, other fsCNA (non-BFB, non-ecDNA) status, *TP53* disruption (at least one gene copy affected), whole-genome duplication (WGD) status (from Paulson et al., 2022), chromothripsis status (Paulson et al., 2022), as well as on-level and windowed histology for each time point and both upper and lower esophageal samples for timepoints TP-1 and TP-2. Maximum histology from any histology biopsy is shown at the bottom of each time-point. Asterisk indicates cancer diagnosis made at next endoscopy since biopsies from the diagnostic ESAD endoscopy were unavailable for CO patient ID 772 and lacked sufficient DNA for CO patient ID 568, so biopsies from the penultimate endoscopy were substituted (occurring 1.44 and 8.16 months after TP-2 for patients 568 and 772, respectively).

**Supplemental Figure 3.**
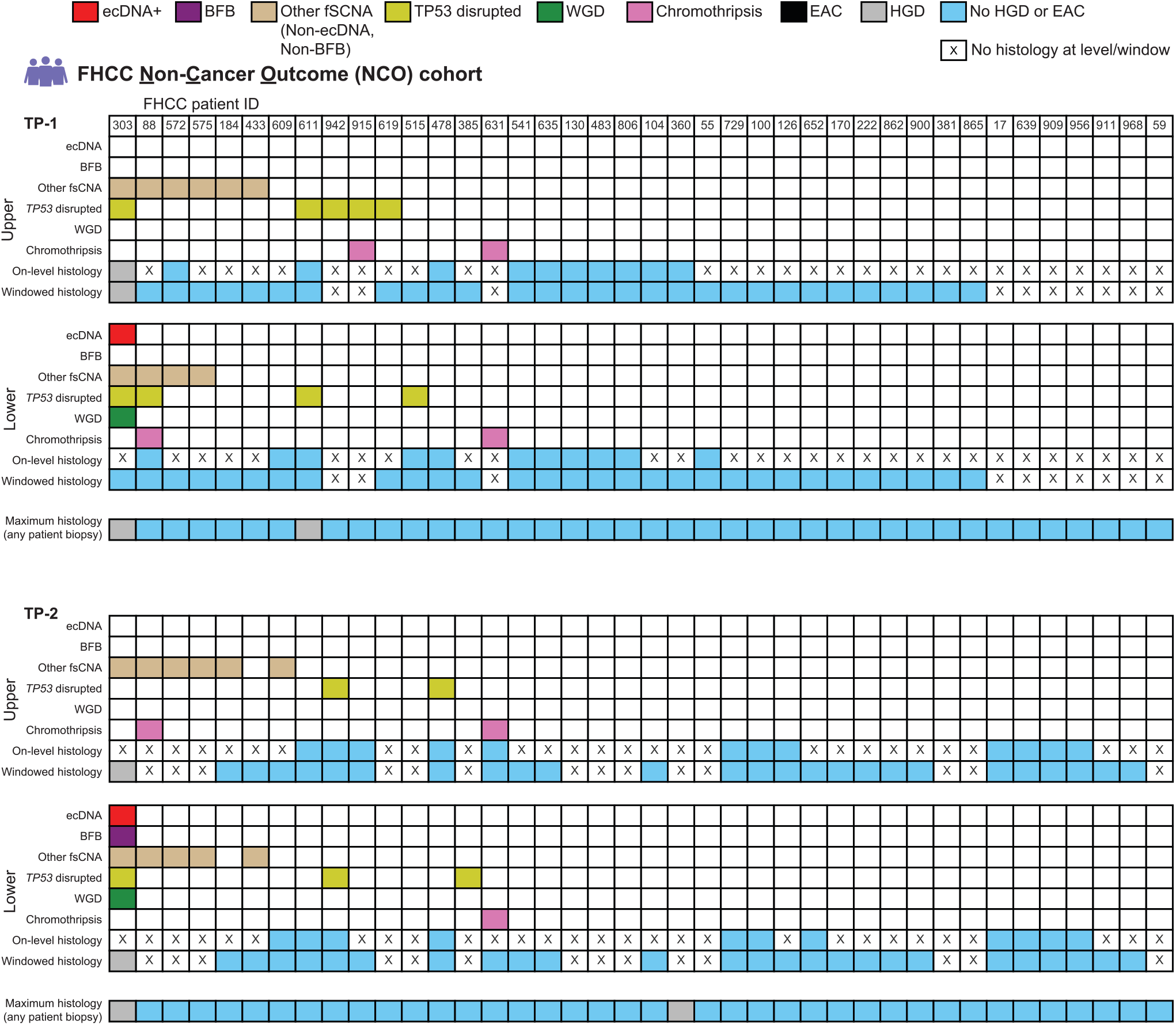
Oncoprint tables of FHCC NCO patient WGS samples encoding ecDNA status, BFB status, other fsCNA (non-BFB, non-ecDNA) status, *TP53* disruption (at least one gene copy affected), whole-genome duplication (WGD) status (from Paulson et al., 2022), chromothripsis status (Paulson et al., 2022), as well as on-level and windowed histology for each time point and both upper and lower esophageal samples for timepoints TP-1 and TP-2. Maximum histology from any histology biopsy is shown at the bottom of each time-point.

**Supplemental Figure 4.**
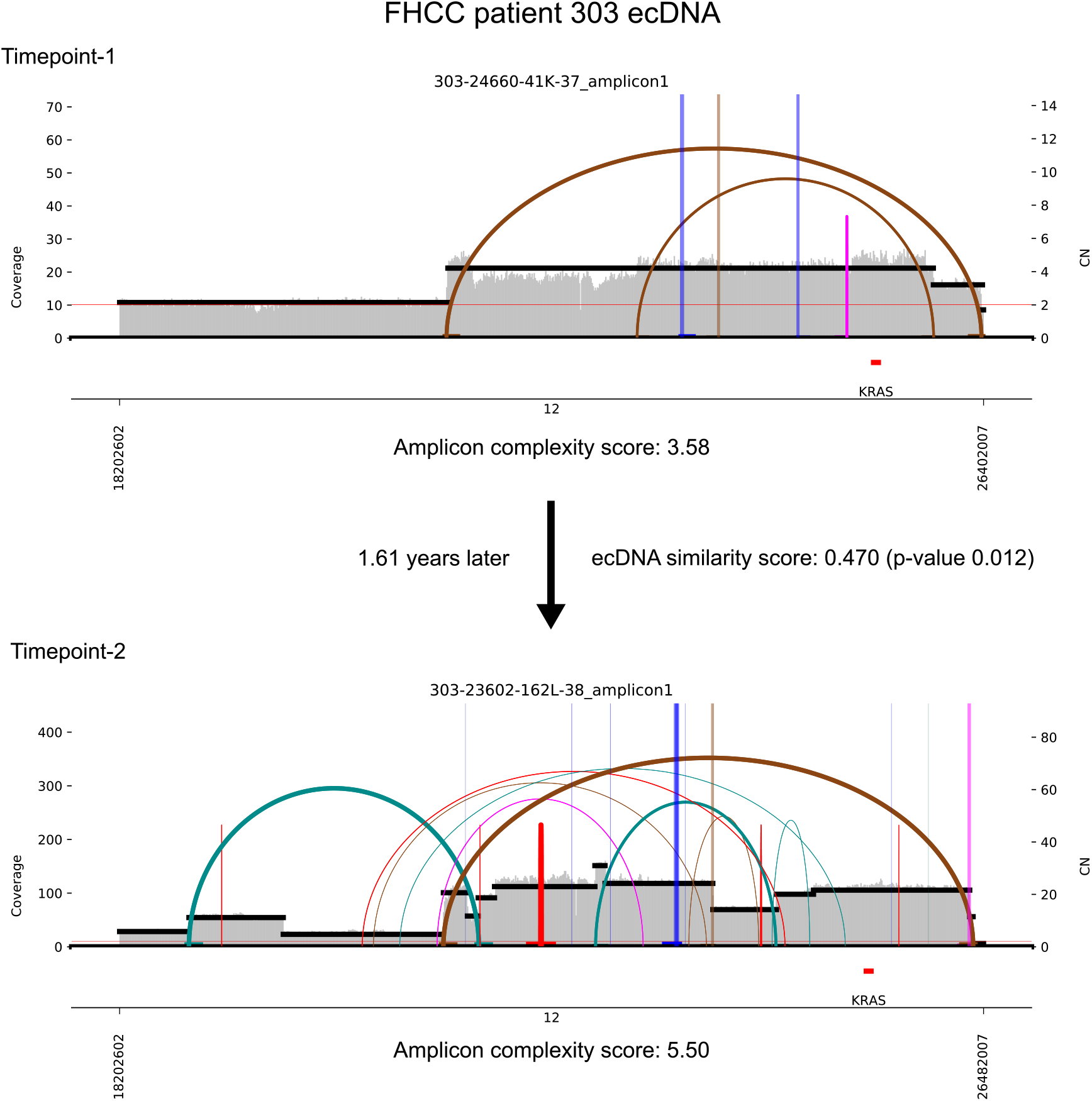
The *KRAS*-bearing ecDNA focal amplification detected in FHCC NCO patient 303 at timepoint TP-1 and timepoint TP-2. Amplicon similarity analysis suggests a common origin of the ecDNA, and ecDNA copy number and complexity increased during the 1.61 years between samples.

**Supplemental Figure 5.**
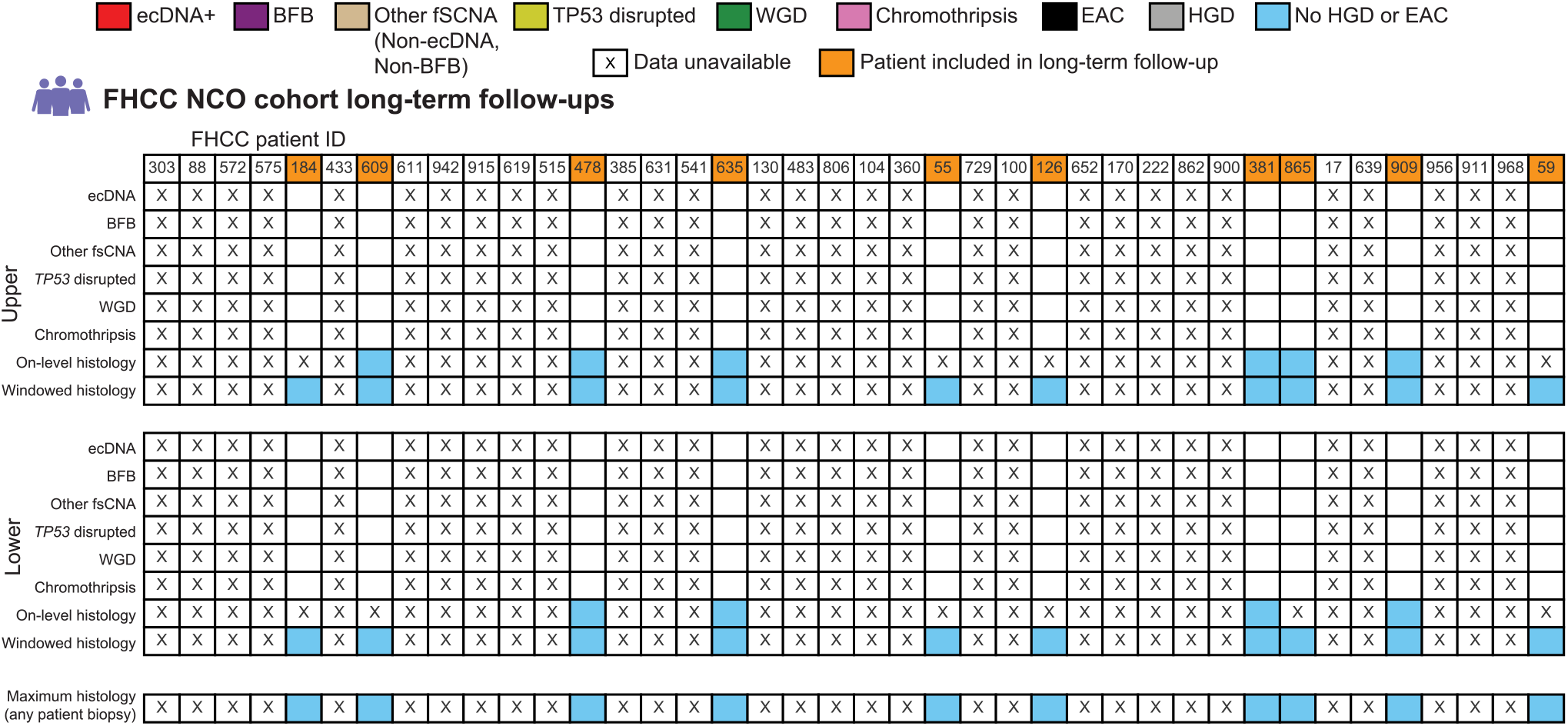
Oncoprint tables of FHCC NCO long-term follow-up patients (collected median 9.6 years after TP-2).

**Supplemental Figure 6.**
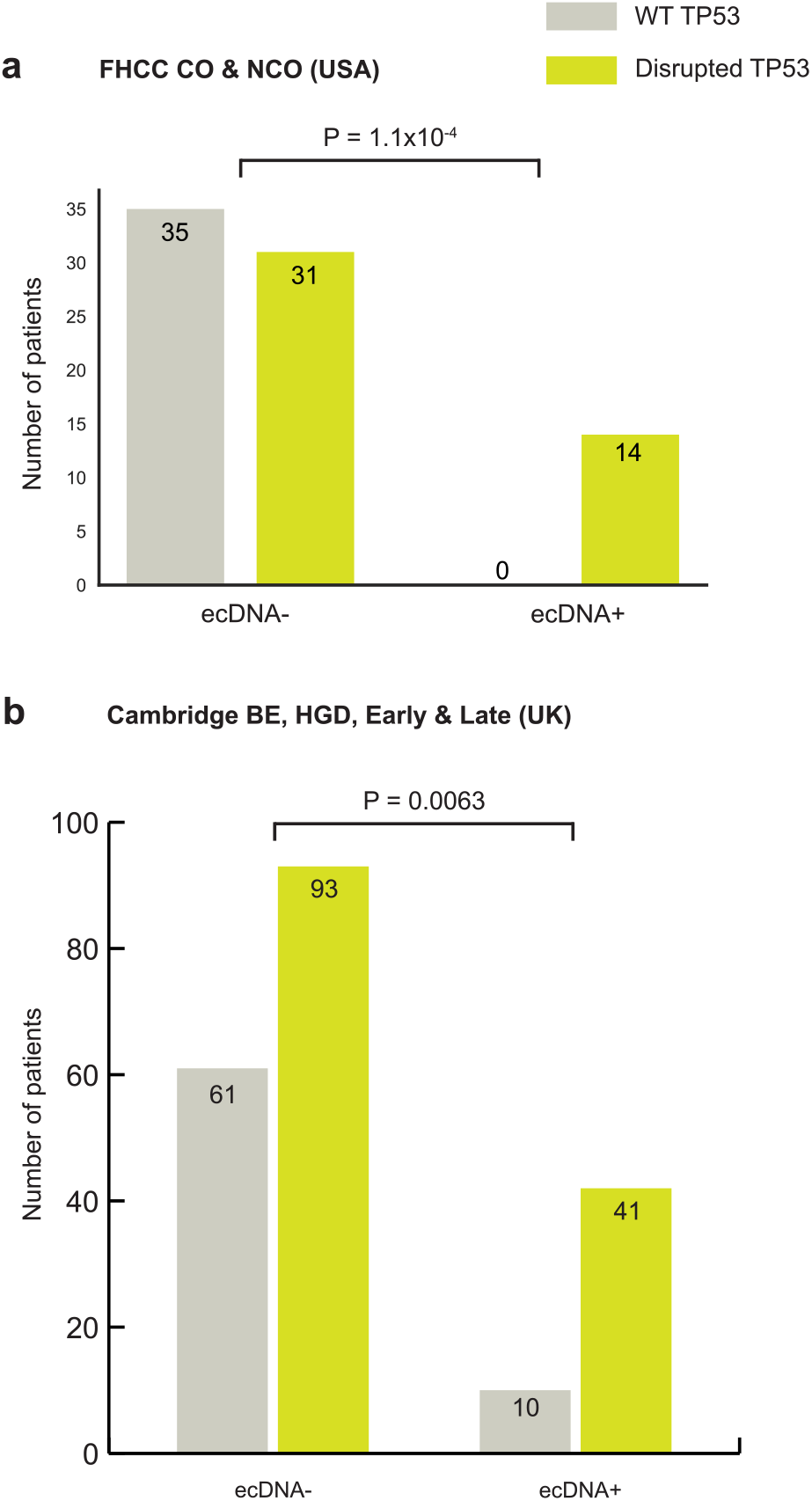
**a)** Association of ecDNA presence and *TP53* status for FHCC cohort patients. **b)** Association of ecDNA presence and *TP53* status for Cambridge cohort patients. Fisher’s exact test, p-values 1.1×10^−4^ and 6.3×10^−3^, respectively for FHCC and Cambridge, H_a_=greater.

**Supplemental Figure 7.**
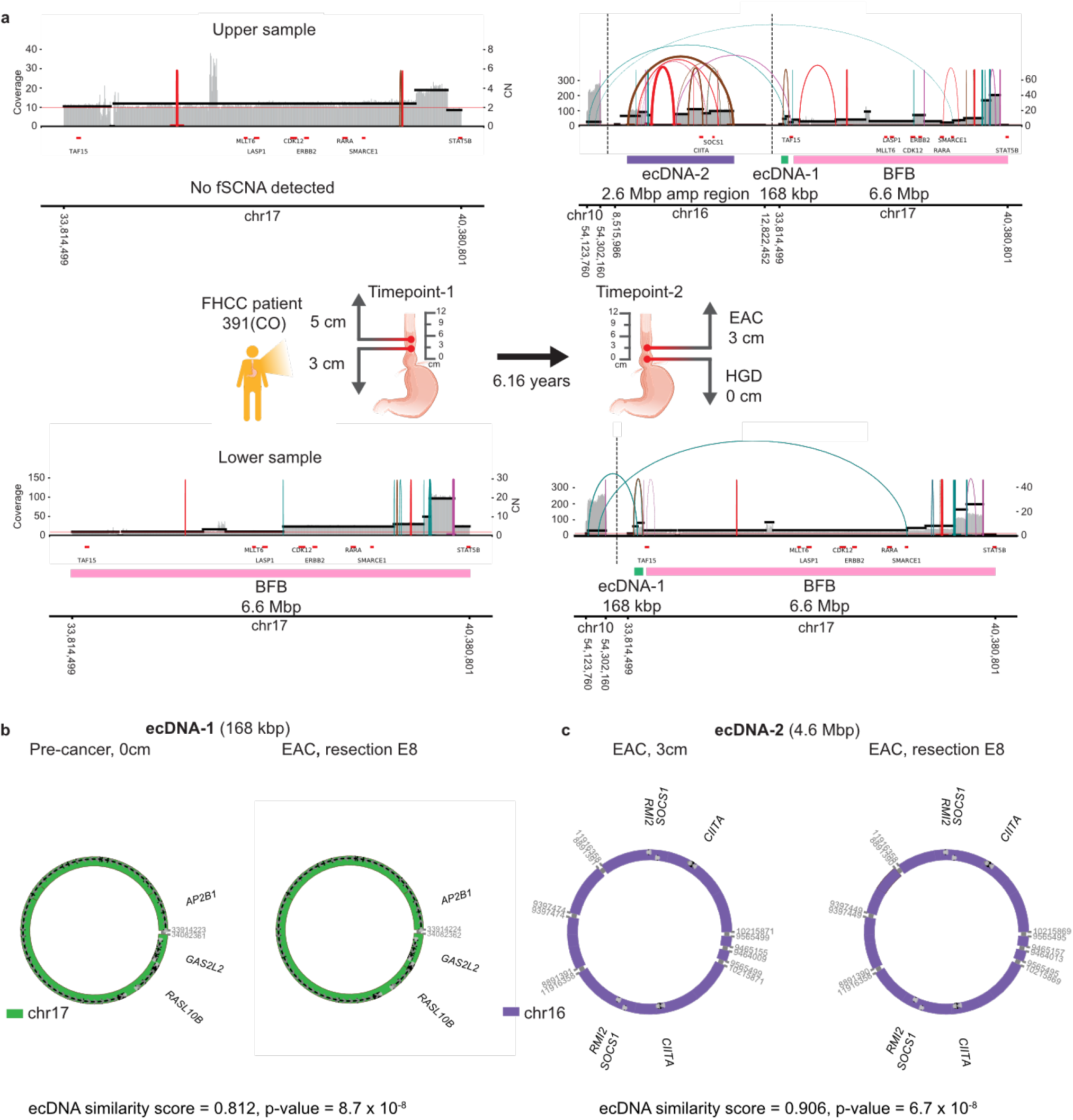
**a)** Patient 391 conserved focal amplification of breakage-fusion-bridge and emergence of ecDNA between timepoints TP-1 and TP-2. **b)** The structure of patient 391’s ecDNA-1, detected in the lower pre-cancer sample from TP-2, where HGD was in the histology window, and an identical structure derived from the adenocarcinoma resection. **c)** The structure of ecDNA-2, detected in the upper sample from TP-2 where EAC was present in the histology window, and an identical structure derived from the adenocarcinoma resection. Amplicon similarity analysis of ecDNA-1 and -2 reveals common origins of the structures.

**Supplemental Figure 8.**
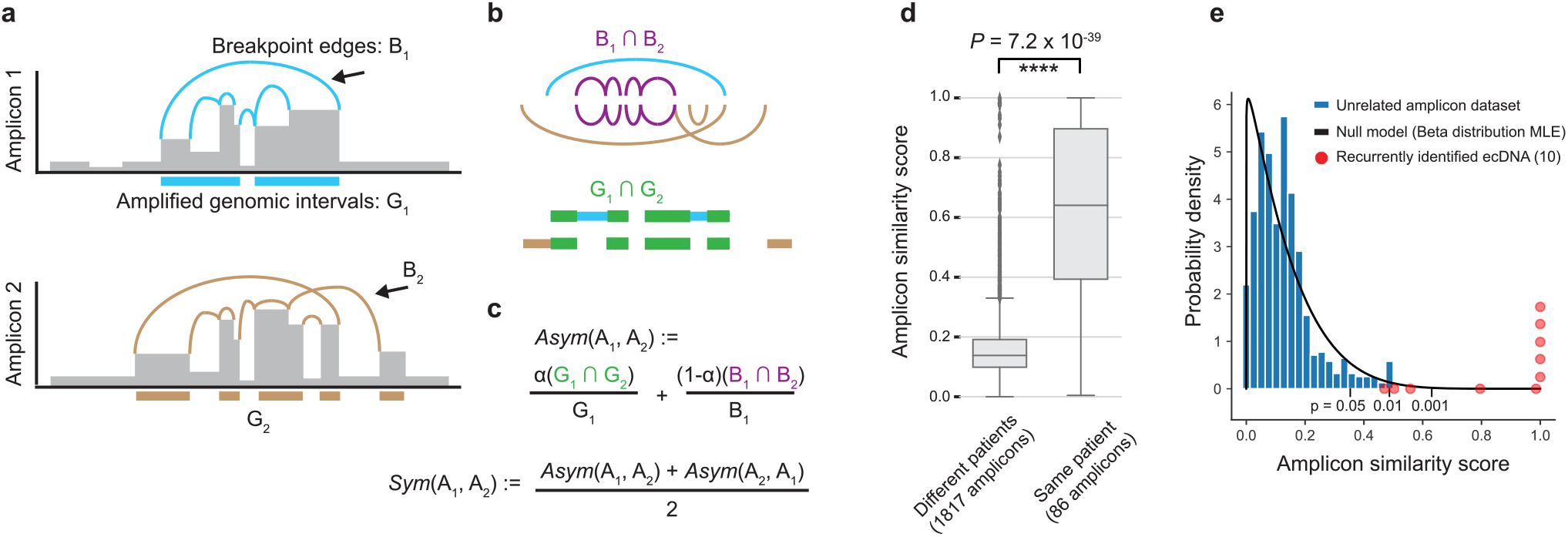
**a)** Cartoon representation of two overlapping focal amplifications (A_1_, A_2_) consisting of a collection of genomic intervals (G_i_) and breakpoints (B_i_). Genomic location is shown on the x-axis and copy number on the y-axis. **b)** (Top) Representation of the relative locations of B_1_, B_2_ and G_1_, G_2_ and the resulting union of those elements, (Bottom) representation of the intersection of the elements in B_1_, B_2_ and G_1_, G_2_ highlighted in purple and green, respectively. **c)** (Top) Definition of the asymmetric similarity score function *Asym* for two overlapping amplicons. (Bottom) Definition of the symmetric similarity score, *Sym* for two overlapping amplicons, which is the average of the asymmetric scores. **d)** The distribution of maximum asymmetric similarity scores for overlapping amplicons derived from different patients (left) and for overlapping amplicons derived from the same patient (right) in FHCC NCO and CO patients shows significantly higher similarity scores for amplicons derived from the same patients (Mann-Whitney U test, p-value=7.2×10^−39^, test statistic=13463.0). **e)** Probability density plot of amplicon similarity scores from a collection of unrelated samples with overlapping focal amplifications (blue), a beta distribution maximum-likelihood estimate of the empirical amplicon similarity score distribution (black), and the similarity scores of overlapping ecDNA amplicons from the same FHCC patients (red).

**Supplemental Figure 9.**
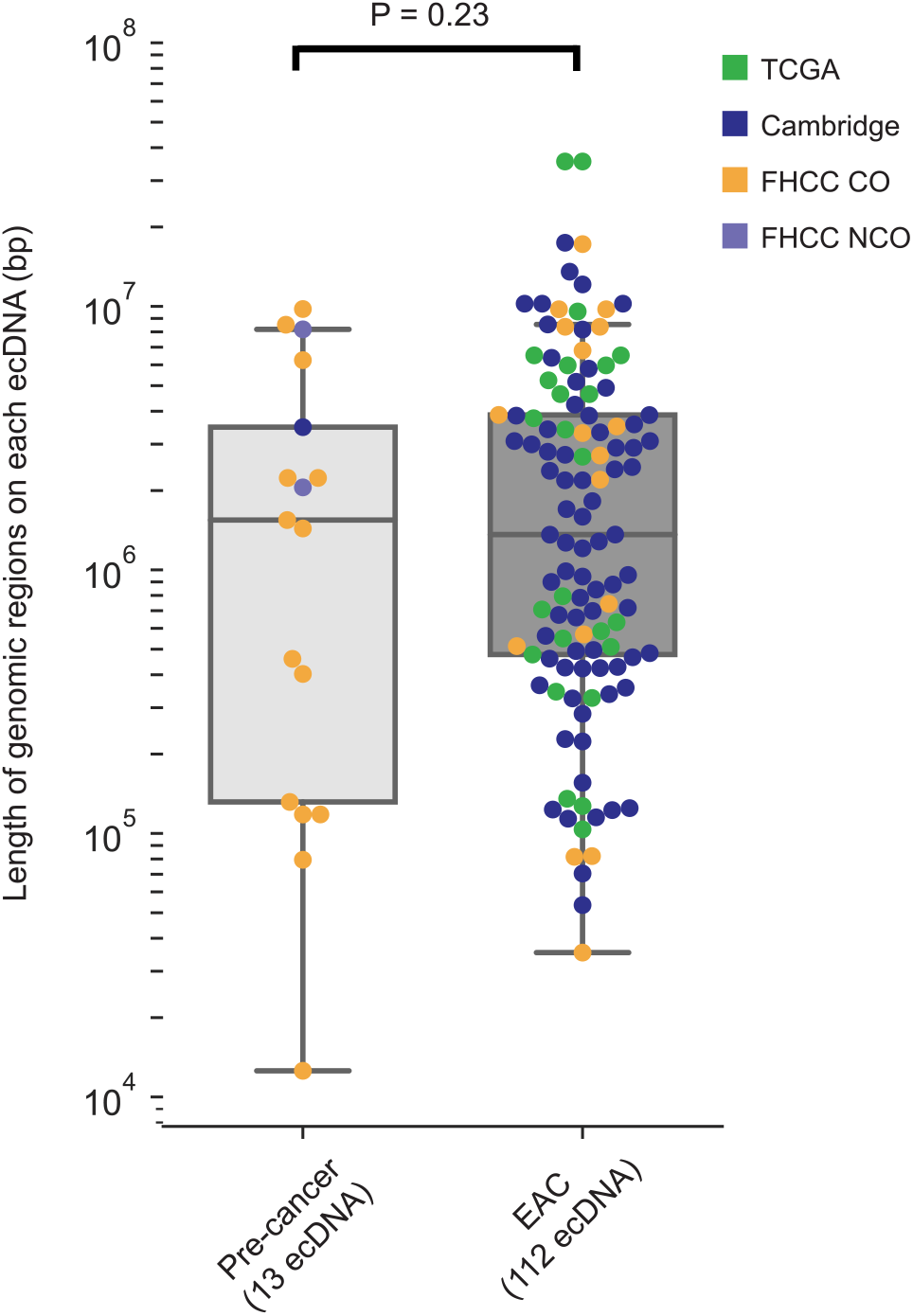
The length of predicted genomic intervals captured on ecDNA, visualized on log10 scale, for each distinct ecDNA in the combined cohorts, segregated by pre-cancer versus EAC shows no significant difference in length of intervals captured on ecDNA between pre-cancer and EAC ecDNA (Mann-Whitney U test, p-value=0.23).

**Supplemental Figure 10.**
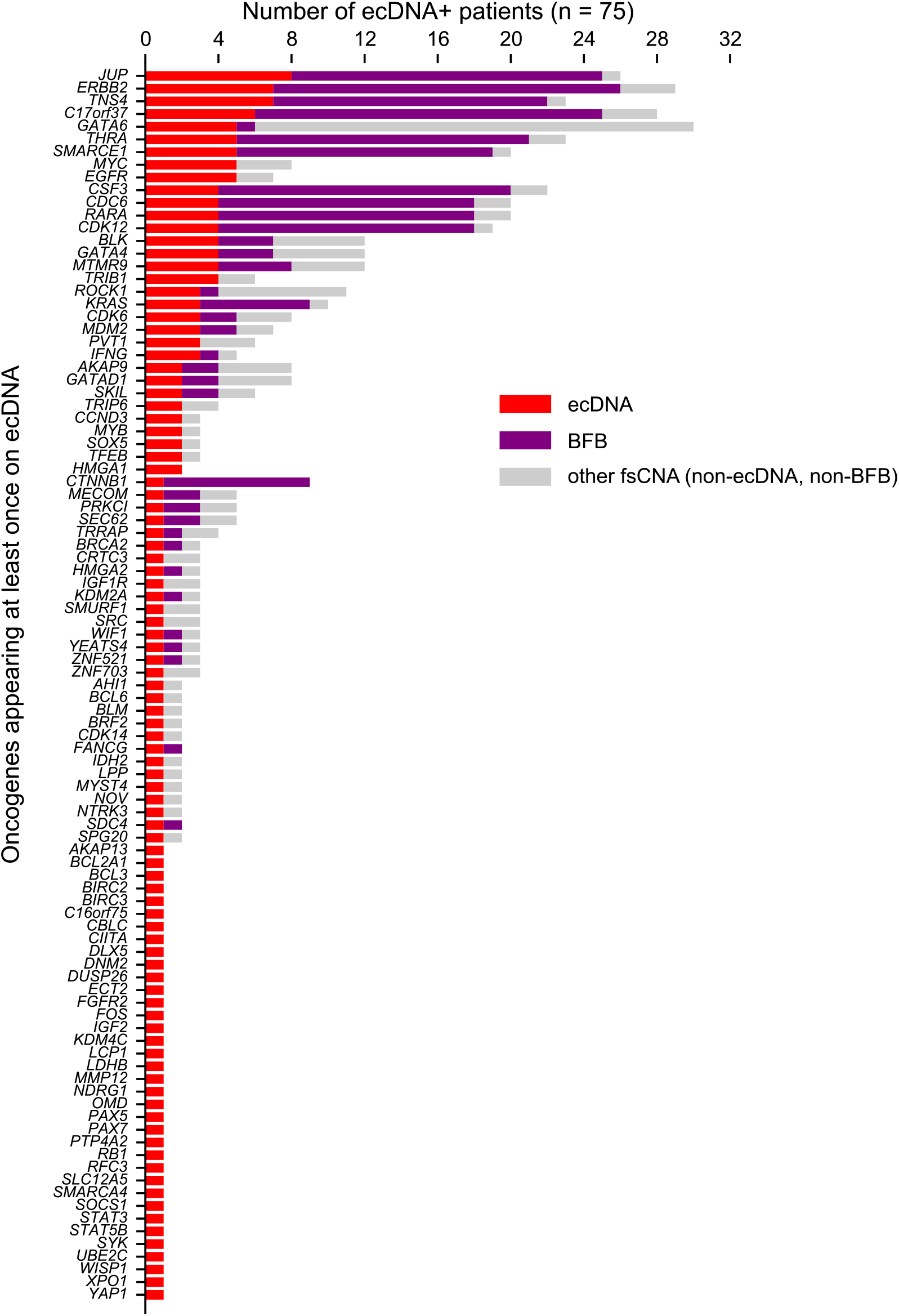
**a)** For oncogenes detected on ecDNA in at least one patient, the number of ecDNA-positive patients having the oncogene listed on ecDNA, and the frequency of that gene on other focal amplifications.

**Supplemental Figure 11.**
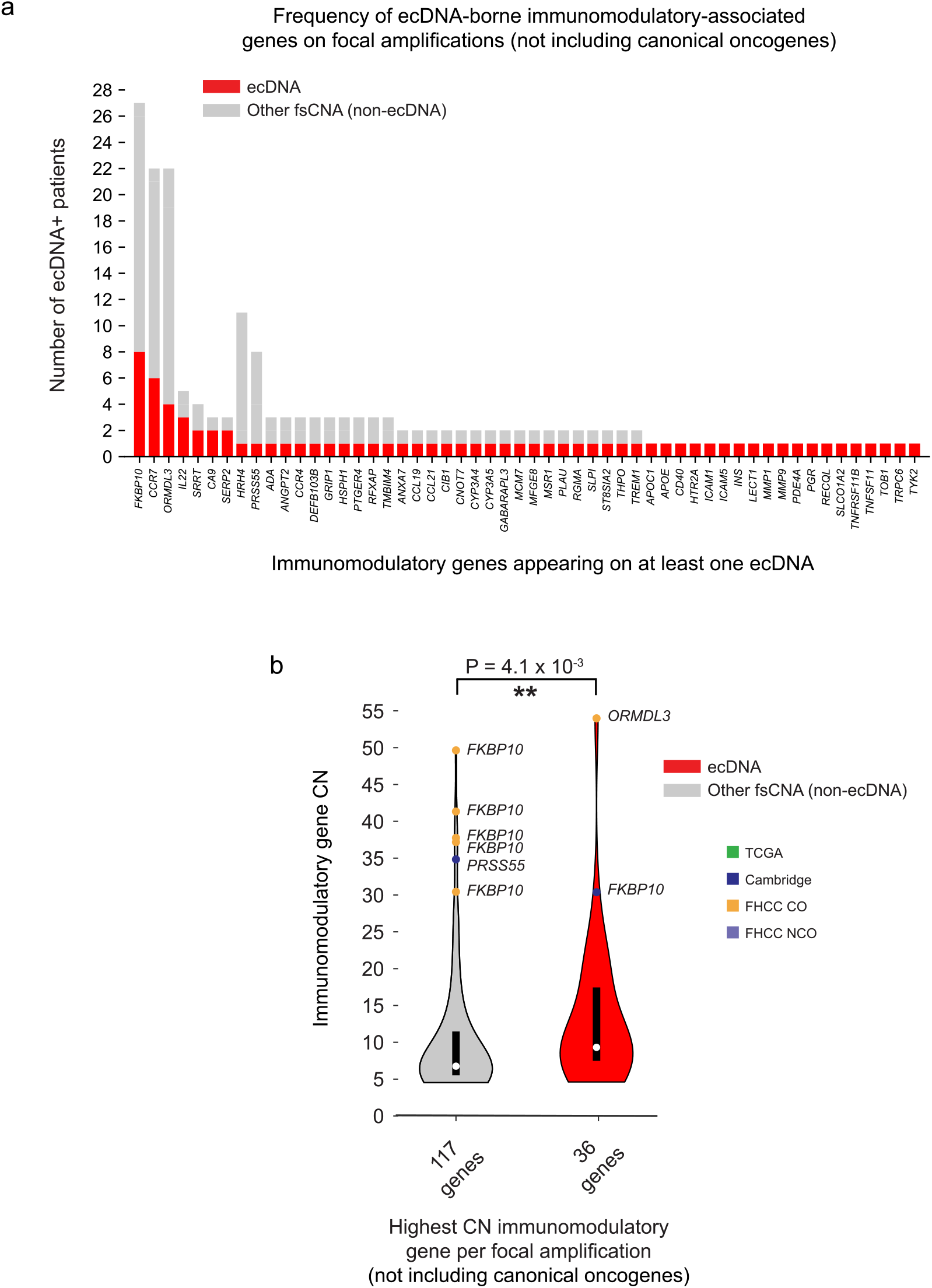
**a)** For immunomodulatory-associated genes detected on ecDNA in at least one patient, the number of ecDNA-positive patients having the gene listed on ecDNA, and the frequency of that gene on other focal amplifications. **b)** Distributions of copy numbers for the highest copy number focally amplified immunomodulatory-associated gene in each unique amplicon which was carried on ecDNA or non-ecDNA fsCNA show significantly higher copy number of immunomodulatory-associated genes on ecDNA versus non-ecDNA fsCNA (Mann-Whitney U test, p-value=4.1×10^−3^, test statistic=1490.0).

**Supplemental Figure 12.**
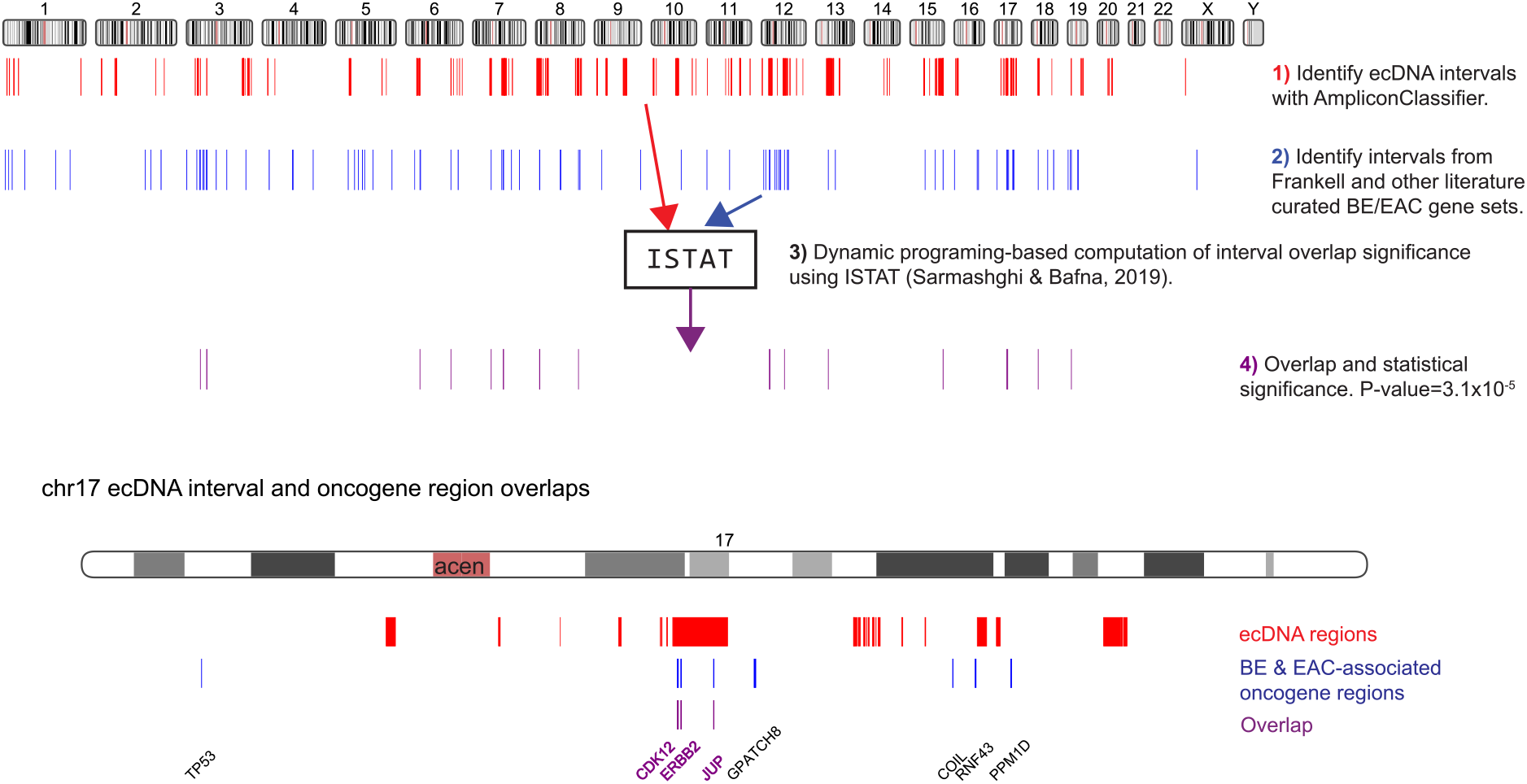
ecDNA and oncogene overlap computation annotated diagram showing how intervals were selected, and the methodology used to compute the overlap statistical significance. ecDNA regions were derived from any ecDNA-positive sample identified in our study. The lower part illustrates the overlap between ecDNA regions and canonical BE- and EAC-associated oncogenes for chr17.

## SUPPLEMENTARY APPENDIX

### Cambridge sample selection

The Cambridge cohort consists of Barrett’s Esophagus (BE) cases with 42 patients having low grade disease, 25 with high grade disease, and 50 early-stage (T1) esophageal adenocarcinoma (EAC). Patients with low grade BE and high grade BE underwent surveillance at Cambridge University Hospitals NHS Trust and consented prospectively to a biomarker and genomic characterization study (Cell Determinants Biomarker, REC no. 01/149, BEST2 REC no. 10/H0308/71). For all samples, strict pathology consensus review was carried out, with 30% of pathological cellularity required for Barrett’s samples and 70% percent for early-stage cancers. BE research samples were collected every 2cm of the BE segment at endoscopy and snap-frozen. A snap frozen section was taken from each BE sample to determine the grade of dysplasia. If more than one grade was present in a sample, it was classified according to the highest grade. In cases which progressed to multiple different disease stages, the highest grade of dysplasia in the case’s follow-up was used for sequencing. Patients in the pre-cancer categories who received prior ablative treatment were excluded. Samples with squamous contamination were excluded.

Early-stage EAC patients were recruited for the EAC International Cancer Genome Consortium (ICGC) study, for which samples were collected through the UK-wide Oesophageal Cancer Classification and Molecular Stratification (OCCAMS, Rec. no. 10-H0305-1) consortium. Ethical approvals for these trials were from the East of England-Cambridge Central Research Ethics Committee. Early-stage EAC samples were prospectively collected as endoscopic biopsies or resection specimens. All tissue samples were snap frozen and blood or normal squamous epithelium (at least 5cm from the tumor) were used as germline reference as previously described^1^.

### Cambridge sequencing data

Sequencing was carried out for cases with an estimated tumour purity of >70% determined by expert pathologist review. Whole genome sequencing by Illumina (100-150bp paired end reads) was carried out with 50-fold coverage for the tumour and 30-fold coverage for the matched germline control. Reads were then aligned with BWA-MEM^2^ to GRCh37 (1000 Genomes Project human_g1k_v37 with decoy sequences hs37d5).

### Cambridge focal amplification detection

Both Cambridge BAM files were aligned to GRCh37 (1000 Genomes Project human_g1k_v37 with decoy sequences hs37d5) using BWA-mem (v0.7.17). Absolute copy number (CN) profiles were generated using ASCAT^3^ (v2.3). Genomic regions with a total CN > 4.5 and interval size > 10kbp were identified, merged, and refined with the amplified_intervals.py script. Each seed region was given to AA separately to improve runtime on each sample. AA was run in the default explore mode to reconstruct amplicon structures and amplicons formed by the same regions were deduplicated based on genomic overlap such that the highest-level classification amplicon was kept (ranked by ecDNA, BFB, complex non-cyclic, and then linear), ties being broken by largest amplicon size.

### TCGA focal amplification detection

We utilized the Dockerized PrepareAA wrapper to detect focal amplifications in the TCGA cohort. The wrapper pipeline for seed detection incorporated CNVKit^4^ (version 0.9.7) run in unpaired mode to detect CNVs. The CNV calls were then provided the amplified_intervals.py script and filtered based on regions having CN > 4.5 and size > 50kbp to produce a set of seed regions. We used AmpliconArchitect^5^ (version 1.2) to infer the architecture of amplicons, The pipeline was run on 20 TCGA-ESCA EAC tumor whole genome sequencing BAMs, aligned to GRCh37, through the Institute for Systems Biology Cancer Genomics Cloud (https://isb-cgc.appspot.com/) which provides a cloud-based platform for TCGA data analysis.

### FHCC sequencing data and annotations

Sequencing data for the FHCC study was previously published in Paulson et al.^6^. All research participants contributing clinical data and biospecimens to this study provided written informed consent, subject to oversight by the Fred Hutchinson Cancer Research Center IRB Committee D (Reg ID 5619). Reads were then aligned with BWA-MEM (version 0.6.2-r126)^2^ to GRCh37 (1000 Genomes Project human_g1k_v37 with decoy sequence hs37d5). BAM files went subsequent indel realignment with GATK IndelRealigner^7^ (version 3.4-0-g7e26428). Chromothripsis calls were derived from Hadi et al.^8^. Genome doubling (WGD) calls were derived from Paulson et al.^6^.

### FHCC cohort focal amplification detection

We utilized the PrepareAA wrapper to detect focal amplifications in the FHCC cohort. The wrapper pipeline for seed detection incorporated CNVKit^4^ (version 0.96) run in tumor-normal mode to call somatic CNVs against the matched normal WGS samples for each patient (when multiple normal samples were available, one was selected arbitrarily). Normal samples also underwent the same pipeline in unpaired mode for standalone CNV detection. The CNV calls were then provided the amplified_intervals.py script and filtered based on regions having CN > 4.3 (4.0 for normals) and size > 50kbp (10kbp for normals) to produce a set of seed regions. The wrapper then invoked AmpliconArchitect (version 1.2) in default mode on the WGS bam files to examine seed regions and profile the architecture of the focal amplifications. The resulting graph and cycles output files were provided to AmpliconClassifier (AC) (version 0.4.5) to produce classifications of the AA amplicons for ecDNA, BFB, complex non-cyclic and linear focal amplifications (Supplementary Appendix - “Amplicon classification and ecDNA detection”). AC also specified bed files corresponding to the classified regions and annotated the identity of genes on the focal amplifications.

### FHCC cohort histology

In the FHCC cohort histology and sequencing biopsies were collected separately. If a sequencing biopsy had a histology biopsy from the same level along the esophagus (measured from the gastroesophageal junction), then it was denoted as having on-level histology. If a sequencing biopsy had a histology biopsy from within +/- 1cm of the same level, it was denoted as having windowed histology. When multiple histology samples could be paired with the sequencing, the histology biopsy with most severe disease state was assigned.

### TP53 disruption analysis

In the FHCC cohort, *TP53* status was determined from Paulson et al.^6^ and we defined *TP53* disruption where either single (+/-) or double (-/-) loss of *TP53* was detected. In brief, for the FHCC cohort, mutations were defined as any moderate-to high-impact SNV or indel as reported by SNPeff^9^. Deletions of at least one exon, or SVs affecting the *TP53* coding sequence or splice sites were also considered to disrupt *TP53*, as were copy number alterations affecting at least half of exonic regions. All alterations were verified manually using IGV^10^ or Partek®. For the Cambridge cohort, *TP53* status was determined by identifying somatic coding variants (missense, frameshift, stop gain or splice site variants), using Strelka^11^ v2.0.15 and Variant Effect Predictor^12^ (VEP) version 78. Disruption was defined as one or more copies of *TP53* being affected by a mutational event.

### Selection of gene lists

Oncogenes were derived from a combination of the ONGene database^13^, as well as BE & EAC driver genes listed from Frankell et al.^1^, Stachler et al.^14^, and Paulson et al.^6^. The complete list is given in Supplemental Table 5. Immunomodulatory genes were derived from the HisgAtlas database^15^. When evaluating the presence of genes on ecDNA, the average gene copy number was required to be 4.5 or higher and the 5’ end intact.

### Amplicon classification and ecDNA detection

We utilized AmpliconClassifier (AC) (version 0.4.9, available at https://github.com/jluebeck/AmpliconClassifier) to perform classification of AA outputs into different types of focal amplifications and to extract coordinates of the genomic regions corresponding to those classifications.

AC takes two primary inputs - the AA breakpoint graph file encoding genomic segment copy numbers and SV breakpoint junctions, as well as the AA cycles file encoding decompositions of the AA graph file into overlapping cyclic and/or non-cyclic paths weighted by the portion of the genomic CN they represent. AmpliconClassifier uses multiple heuristics to perform the classifications. First AC filters the paths <10kbp, paths which significantly overlap low-complexity or repetitive regions, paths which overlap regions of the genome never exceeding CN 4.5 (not focally amplified), or which have a decomposed CN < δ (too low-frequency relative to other decompositions for reliable classification as focal amplifications). The decomposed CN (*c*_*p*_) threshold, δ, for a path *p*, having a maximum genomic CN of *m*_*p*_ is defined as

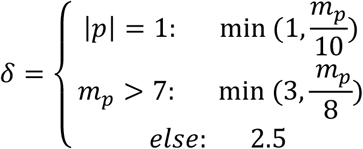

For each remaining path, AC computes a length-weighted CN, called *W*, which is the product of the length of the path (in kbp) and the decomposed path’s assigned copy number.

AC first assess non-filtered paths for the presence of BFB cycles using heuristics determined from manual examination of BFB-like focal amplifications in the FHCC cohort and focal amplifications in previous studies^5,16^. AC computes the fraction of breakpoint graph discordant edges which are foldback, *f*, – i.e., inverted orientation having a genomic distance < 25kbp. AC then identifies decomposed paths containing foldback junctions between segments, and using all paths computes the set of consecutive segment pairs in the paths where the two boundaries of the segments together form a foldback junction. Each segment pair is assigned its own weight equal to the decomposed copy count of the path. If the proportion of BFB-like segment pairs over all segment pairs in all paths is less than 0.295, then the amplicon is not considered to contain a BFB. Furthermore, if the total weights of pairs which are “distal” (not foldback and > 5kbp jump between endpoints) divided by the total weight of all pairs is greater than 0.5, the amplicon is not considered to contain BFB. Lastly, if the total decomposed CN of all pairs is < 1.5, or if the total number of foldback segment pairs is < 3, or *f* < 0.25, or the decomposed CN weight of all BFB-like paths divided by the CN weight of all paths < 0.6, or the maximum genomic copy number of any region in the candidate BFB region is < 4, the amplicon is not considered to contain a BFB. If the amplicon has not failed any of these criteria, a BFB-positive status is assigned, and the BFB-like cycles (decomposed paths with a BFB foldback) are put into a set and kept separate from additional fsCNA detection inside the amplicon region.

Next, AC assess non-filtered, non-BFB paths for the presence of ecDNA cycles. If there is any cyclic path with decomposed CN > 5 and length > 100kbp, an ecDNA-positive status is assigned. If the total fraction of length-weighted CN, *W*, assigned to cycles exceeds 12% of the total length-weighted CN in the cycles file and more than 10kbp are inside the filtered cyclic paths, an ecDNA-positive status is assigned. Lastly, if the total length of complex cycles (cyclic paths with interior rearrangements > 5kbp) exceeds 50kbp and the region has CN > 4.5 an ecDNA-positive status is assigned. The ecDNA-like cyclic paths are then stored for subsequent analysis, including reporting of the genomic coordinates as a bed file and annotation of genes.

If the amplicon is not classified as BFB-positive and/or ecDNA-positive, and has paths consistent with focal amplification, then two other classifications are checked. If the fraction of *W* assigned to non-cyclic paths with rearrangements > 5kbp plus *W* assigned to cyclic paths is greater than 0.3 of total *W* in all paths, a complex non-cyclic label is assigned. If the ratio of *W* assigned to non-cyclic paths without rearrangements to *W* assigned to non-amplified paths is greater than 0.25, then the path is labeled complex non-cyclic if the breakpoint graph has > 4 discordant edges in amplified regions, otherwise a linear amplification label is assigned. If not resolved by these heuristics, the path type with the highest fraction of *W* is assigned.

### Amplicon similarity score

We compared overlapping focal amplifications to quantify amplicon similarity by quantifying the relative amounts of shared overlap in genomic coordinates and in SV breakpoint location. These calculations are implemented into the amplicon_similarity.py script, available in the AmpliconClassifier repository (https://github.com/jluebeck/AmpliconClassifier).

We defined symmetric and asymmetric amplicon similarity scores combining information from both the genomic interval overlap and the shared breakpoint junctions. An amplicon is defined as a collection of breakpoints (*B*), and genomic segments (*G*). Genomic overlap was evaluated on the basis of the number overlapping base-level coordinates in two intervals. Breakpoints were considered to be shared if the total distance between the two endpoints of each junction was in total measured to be less than *d* (default = 250bp). That is, for two breakpoints *x* and *y* with sorted endpoints (*x*_*1*_,*x*_*2*_) and (*y*_*1*_,*y*_*2*_), respectively, they must satisfy

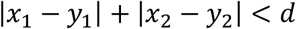

The asymmetric amplicon similarity score between two amplicons A_1_ and A_2_ we defined

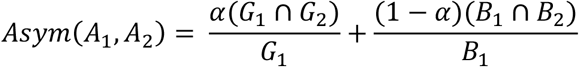

and similarly, the similarity of A_2_ to A_1_ is

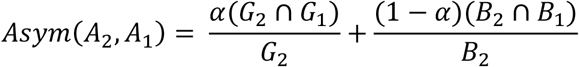

Where α is set to 0.25 by default. We then define a symmetric amplicon similarity score which is the average of the two asymmetric scores

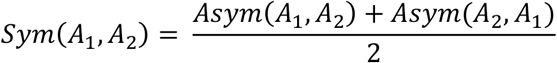

We computed symmetric amplicon similarity scores for a panel of amplicons from unrelated origins derived from sequencing data published in Deshpande et al.^5^, deCarvalho et al.^17^, as well as Steele et al.^18^ and Moody et al.^19^ (using AA amplicons reported in Bergstrom et al.^20^), and the amplicons from unrelated patients in the FHCC cohort. We used the resulting distribution of 719 similarity scores for overlapping amplicons as a background null distribution. We computed the percentile of each new amplicon similarity score in this null distribution to quantify its similarity against the panel of overlapping amplicons from unrelated origins.

We also fit a beta distribution to the empirical null symmetric similarity score distribution, using a maximum likelihood estimation approach to fit the parameters of the model. The beta distribution was selected as it provides support on the interval [0, 1], provides a higher degree of flexibility in fitting various distributions given the two shape parameters, and enables a better estimation of small p-values than the empirical dataset. We performed negative log likelihood minimization using the SciPy^21^ (version 0.19.1) *fmin* function with initial parameter estimates (1.5, 10), and convergence occurred in 38 iterations.

As AmpliconArchitect may include flanking regions which are not focally amplified as part of the amplification itself, the amplicon similarity script filters from the calculation regions that are not focally amplified (CN < 4.5 default), SVs which join two elements less than 2500bp away, and it redundantly filters regions that are also present in the low-complexity or low-mappability database used by AmpliconArchitect.

### Amplicon complexity score

AmpliconArchitect outputs a collection of (cyclic and/or non-cyclic) paths in the CN-aware breakpoint graph representing an approximate optimal balanced CN flow in the graph. As a result, non-trivial graphs may be decomposed into multiple paths, each having some copy-number assigned to the path, constrained by the total amount of CN flow available in the graph. Each path has a copy number *c*, and a length in kilobase pairs, *s*. The total length-weighted copy number of all decomposed paths we call *T*, and is given by

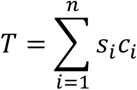

Where *c*_*i*_ (*s*_*i*_) is the copy number (length) of the *i*-th path. The values of *c*_*i*_ are pre-sorted in descending order for increasing *i*. For the decomposed paths of each amplicon graph, *G*, we computed a vector representing the fraction of total CN captured by each of the *n* decompositions. We denote this sorted collection as,

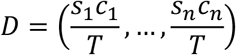

We noted that there may be many low-weight CN paths, representing non- or weakly-amplified paths extracted from the graph, and thus we defined a “residual”, measured against the first percentile, *p*, (default = 80%) of weighted CN explained. We first define an index *j*, where *j* is the largest value such that

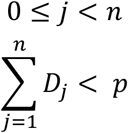

This implies that *j*+1 represents the first index such that sum of the first *j*+1 entries is equal to or exceeds *p*. The residual, *ϵ*, we defined as the weighted CN fractions above the first j+2 entries, is then given by

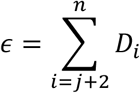

We then defined an amplicon complexity score function *H(ϵ, D, k)*, represented by the sum of entropies from the residual, the non-residual, and the total number of segments in the breakpoint graph, *k*.

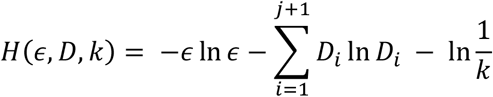

